# Insight into motility-dependent pathogenicity of the zoonotic spirochete *Leptospira*

**DOI:** 10.1101/2020.02.11.944587

**Authors:** Jun Xu, Nobuo Koizumi, Shuichi Nakamura

**Affiliations:** Department of Animal Microbiology, Graduate School of Agricultural Science, Tohoku University, Sendai, Miyagi, Japan; Department of Bacteriology I, National Institute of Infectious Diseases, Shinjuku-ku, Tokyo, Japan; Department of Applied Physics, Graduate School of Engineering, Tohoku University, Sendai, Miyagi, Japan

## Abstract

Bacterial motility is crucial for many pathogenic species in the process of invasion and/or dissemination. The spirochete bacteria *Leptospira* spp. cause symptoms, such as hemorrhage, jaundice, and nephritis, in diverse mammals including humans. Although loss-of-motility attenuate the spirochete, the mechanism of the motility-dependent pathogenicity is unknown. Here, focusing on that *Leptospira* spp. swim in liquid and crawl on solid surfaces, we investigated the spirochetal dynamics on the host tissues by infecting cultured kidney cells from various species with pathogenic and nonpathogenic leptospires. We found that, in the case of the pathogenic leptospires, a larger fraction of bacteria attached to the host cells and persistently traveled long distances using the crawling mechanism. Our results associate the kinetics and kinematic features of the spirochetal pathogens with their virulence.

**One Sentence Summary:** Adhesivity and crawling motility over host tissue surfaces are closely related to the pathogenicity of a zoonotic spirochete.

## Introduction

Many bacteria utilize motility to explore environments for survival and prosperity. For pathogenic species, the motility is a virulence factor, harming the host animal or plant health(*6*). For example, *Helicobacter pylori* requires motility to migrate towards the epithelial tissue in the stomach(*7*), and motility and chemotaxis are key factors that guide host invasion in different *Salmonella* serovars (*8, 9*). Movement and adhesion of the Lyme disease spirochete *Borrelia burgdorferi* in blood vessels are thought to be important during the process of host cell invasion(*10*). In the enteric pathogens, such as the enteropathogenic *Escherichia coli* (EPEC), the motility machinery flagella are also important for adhesion to the host intestinal epithelium(*11*).

In this study, we address the association of bacterial motility with pathogenicity in the worldwide zoonosis leptospirosis. The causative agent of leptospirosis *Leptospira* spp. are Gram-negative bacteria belonging to the phylum Spirochaetes. The genus *Leptospira* is comprised of pathogenic, intermediate, and nonpathogenic species, which are classified into over 300 serovars defined based on the structural diversity of lipopolysaccharide (LPS)(*2, 12*). The pathogenic species affect various mammalian hosts such as livestock (cattle, pigs, horses, and others), companion animals (dogs and others), and humans, causing severe symptoms, such as hemorrhage, jaundice, and nephritis in some host-serovar pairs(*2–4*). The leptospires can be maintained in the renal tubules of recovered animals or reservoir hosts, and the urinary shedding of leptospires to the environment leads to infection in humans and other animals through contact with contaminated soil or water. Although the pathogenic mechanism of leptospirosis is not well elucidated, in addition to the pathogen-specific proteins such as Loa22(*13*), their motility using two periplasmic flagella (PFs) beneath the outer cell membrane (Fig. 1A) is known to be somehow involved in the infection and pathogenicity based on studies in animal models that have shown attenuation of the spirochetes by loss-of-motility(*14, 15*).

**Fig. 1.**
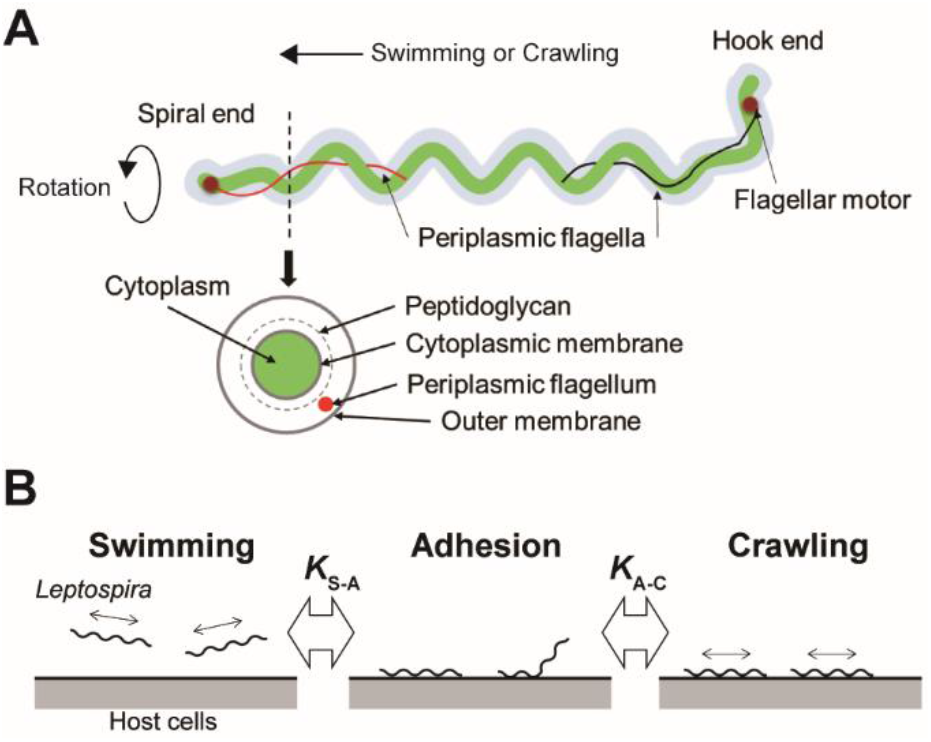
Structure of *Leptospira* and working model. (A) Schematic diagram of the *Leptospira* cell structure. (B) A three-state kinetic model assuming a transition between “swimming” (floating above cell layers without physical contact to the cells) and “adhesion” (attachment to the cell layer without migration), and between “adhesion” and “crawling” (attachment to the cell layer and movement over surfaces), with *K*_S-A_ and *K*_A-C_ represent the equilibrium constants of each transition, respectively.

The rotation of the PFs in *Leptospira* gyrates both ends of the cell body and rotates the coiled cell body, allowing the spirochete not only to swim in fluids but also to crawl over solid surfaces(*16, 17*). Adherence and entry of pathogenic leptospires in the conjunctival epithelium(*4*) and in the paracellular routes of hepatocytes(*18*) were previously observed using scanning electron microscopy, suggesting adhesion to the host tissue surfaces and subsequent crawling of pathogenic leptospires. To verify this hypothesis, assuming the transition of leptospires between swimming and adhesion states and between adhesion and crawling states in the equilibrium (Fig. 1B), we investigated the adhesion and crawling motility of *Leptospira* on the cultured kidney cells of various mammalian species.

## Results

### Steady-state analysis of *Leptospira* on the kidney cells

We infected cultured kidney cells from six different host species (rat, dog, monkey, mouse, cow, and human) with three *Leptospira* strains (the pathogenic *L. interrogans* serovars Icterohaemorrhagiae and Manilae, and the nonpathogenic *L. biflexa* serovar Patoc) expressing green fluorescent protein (GFP) within a chamber slide (Fig. 2A). We observed the *Leptospira* cells by epi-fluorescent microscopy (Fig. 2B) and measured the fractions of swimming [S], adhered [A], and crawling [C] bacteria on the kidney cells (Fig. 2C). Figs. 2D-E show that almost half of the pathogenic population transited from the swimming state to the adhesion and crawling states, whereas >75% non-pathogenic leptospires remained in the swimming state. We calculated the equilibrium constant between swimming and adhesion states (*K*_S-A_) and between adhesion and crawling states (*K*_A-C_) from the cell fractions using *K*_S-A_ = [A] / [S] and *K*_A-C_= [C] / [A], respectively. The pathogenic leptospires had a significantly larger *K*_A-C_ in comparison with the nonpathogenic strain (*P* < 0.05); *K*_S-A_ did not seem to correlate to virulence (Fig. 2F). These thermodynamic parameters suggest that the biased transition from adhesion to crawling would be responsible for the virulence of *Leptospira* (Fig. 2G).

**Fig. 2.**
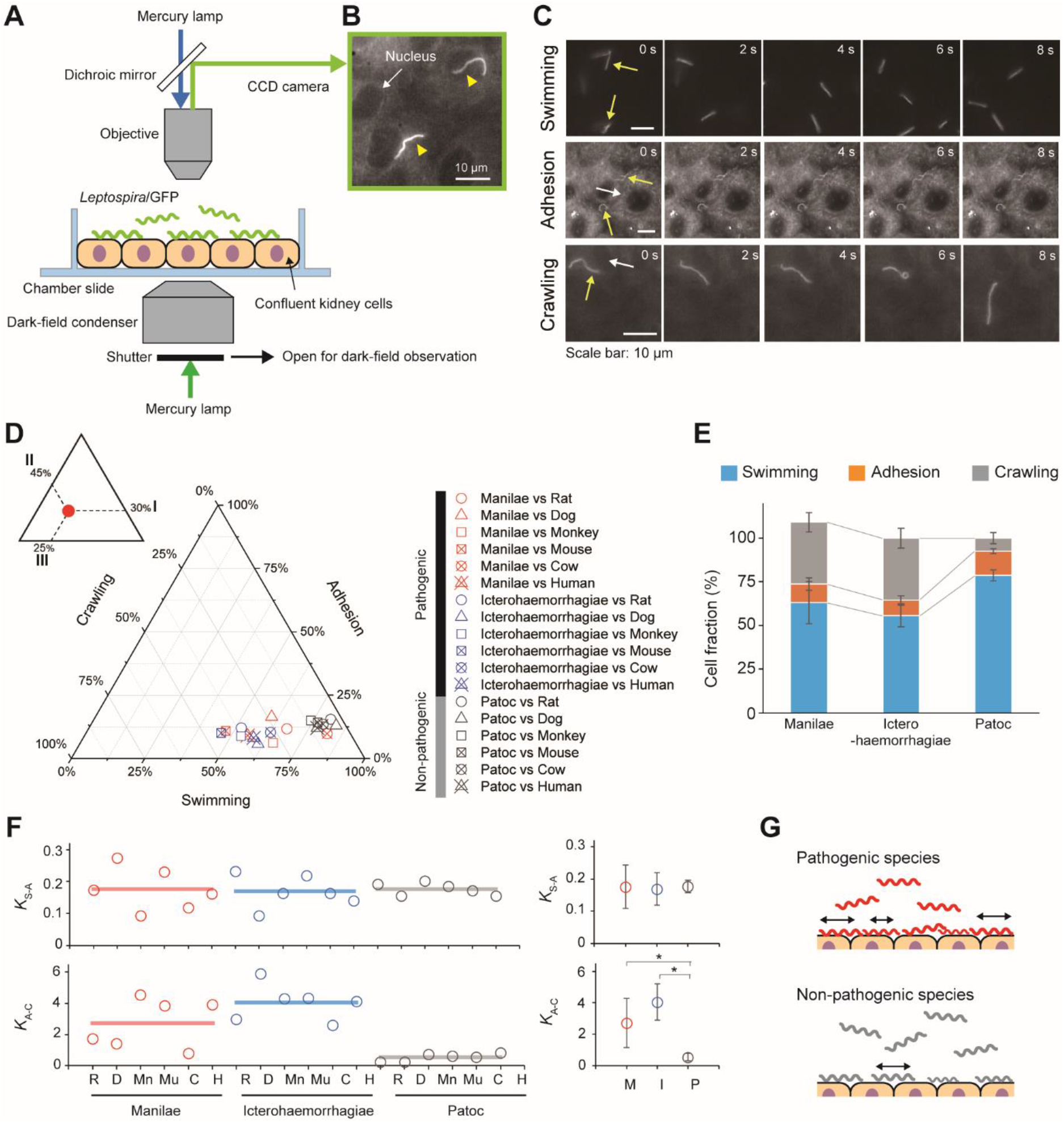
Steady-state motility and adhesion of *Leptospira* cells on kidney cells. (A) Schematic of the motility assay on the cultured kidney cells by epi-fluorescent microscopy. Fully confluent kidney cells from either animals or humans were cultured in an observation chamber, and *Leptospira* cells that constitutively expressed GFP were added to the cultures. (B) Example of an epi-fluorescence image of the *L. interrogans* serovar Icterohaemorrhagiae, as indicated by the yellow triangles, on the rat kidney cell line, NRK-52E. The nucleus of the kidney cell is shown by the white arrow (Movie S1). (C) Image sequences of swimming (top), adhered (middle), and crawling (bottom) leptospiral cells. Yellow and white arrows indicate *Leptospira* cells and kidney cells, respectively. No kidney cells were observed due to out of focus during the measurement of *Leptospira* cells swimming in liquid media. (D) Ternary plot of the cell fractions in a state of swimming, adhesion, or crawling. The inset schematically explains how to read the ternary plot using an example plot with 30% for I, 45% for II, and 25% for III. Legend is shown to the right of the ternary plot, and M, I and P indicate the *L. interrogans* serovar Manilae, *L. interrogans* serovar Icterohaemorrhagiae, and *L. biflexa* serovar Patoc, respectively. Each data point represents the mean of triplicate experiments and ~ 2,400 leptospiral cells were measured per host-serovar pair. (E) Average values of the cell fractions. Error bars are the standard deviation. (F) The host-bacterium dependence of the equilibrium constants *K*_S-A_ and *K*_A-C_, calculated from the data shown in **D**; rat (R), dog (D), monkey (Mn), mouse (Mu), cow (C), and human (H). The averaged values determined for each bacterial strain are shown by horizontal lines and are plotted with the standard deviation in the right; *L. interrogans* Manilae (M), *L. interrogans* Icterohaemorrhagiae (I), and *L. biflexa* Patoc I (P). The statistical analysis was performed by Mann-Whitney U test (* *P* < 0.05). (G) Schematic explanation of the kinetic difference between pathogenic and non-pathogenic leptospires.

### Crawling motility

Taking the results of the steady-state analysis, we focused on the crawling motility of individual *Leptospira* cells on the kidney cells. Although the crawling speed varied among the measured host-bacterium pairs, *L. interrogans* serovar Icterohaemorrhagiae showed significantly faster speed than the others, indicating the species/serovar dependence of the crawling ability (Fig. 3A). On the other hand, there is no difference in crawling speed between *L. interrogans* serovar Manilae and *L. biflexa* serovar Patoc, suggesting that the crawling speed itself is not related to leptospiral virulence. Meanwhile, we observed that some leptospiral cells attached to the kidney cells and moved smoothly for periods, migrating over long-distances (upper panels of Figs. 3B-C; Movie S2), whereas others frequently reversed the crawling direction (lower panels of Figs. 3B-C; Movie S3). Cell movements with reversals are considered to be diffusive and thus can be evaluated by plotting the mean square displacement (MSD) against time, a general methodology for diffusion (Brownian motion) analysis(*19*). The MSD of simple diffusion without directivity is proportional to time, and therefore double-logarithmic MSD plots from such non-directional diffusion represent slopes of ~1, whereas those from directive movements show MSD slopes of ~2, representing the relatively long distance traveled by the cells (Fig. S1). Double-logarithmic MSD plots obtained from each individual leptospires showed a wide range of MSD slopes (example data are shown in Fig. 3D) and differed for each *host-Leptospira* pair (Fig. 3E left and Fig. S2). The nonpathogenic strain showed the slope of ~1, while the pathogenic strains had significantly larger slopes that denote directive motion (Fig. 3E right). Thus, concerning the crawling motility, directivity and persistency rather than speed could be crucial for virulence.

**Fig. 3.**
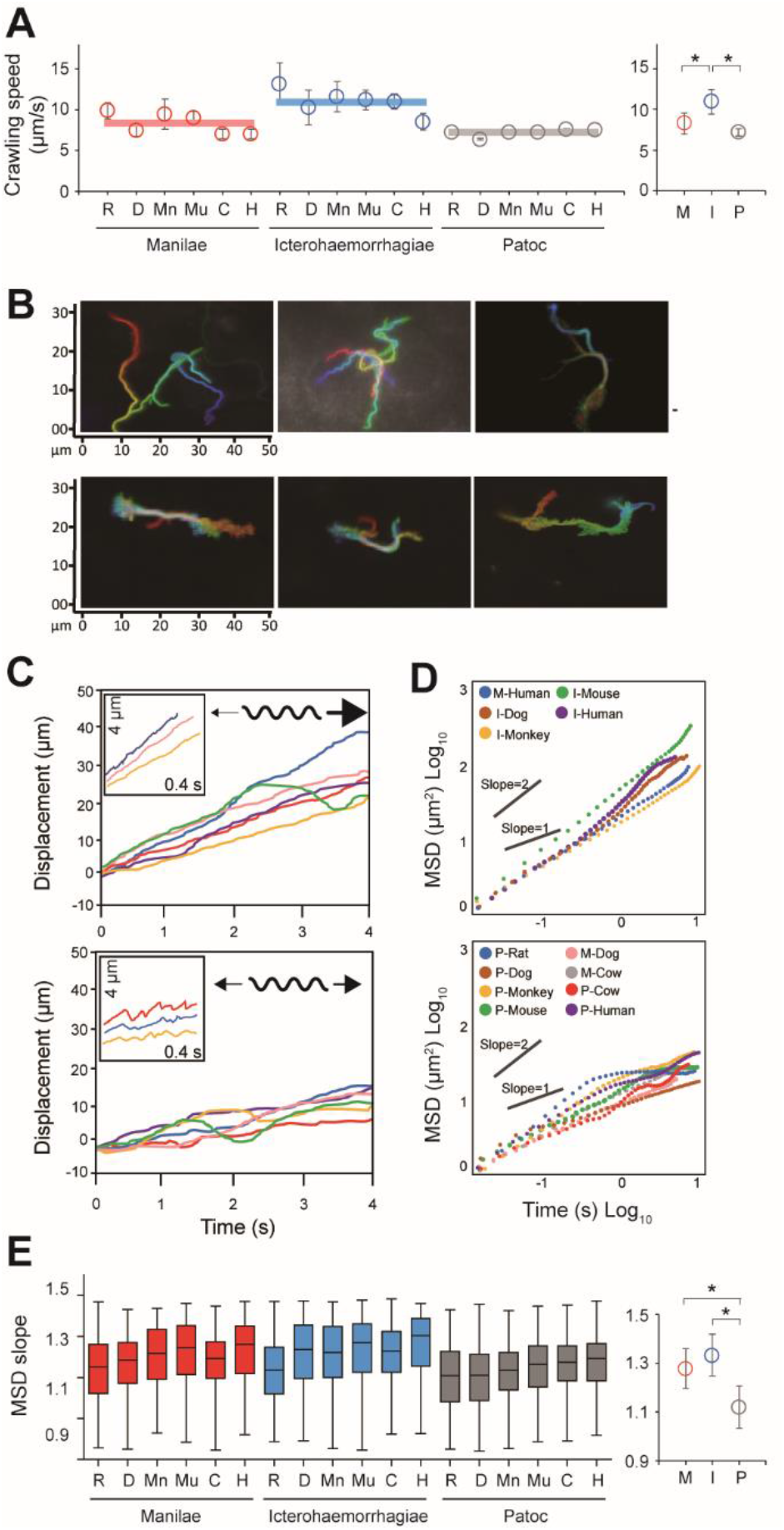
Analysis of the crawling movement of *Leptospira* on kidney cell layers. (A) Average crawling speeds determined for each host-bacterium pair. More than 90 bacteria were measured for each pair. The averaged values calculated for each bacterial strain are shown by horizontal lines (left) and are plotted with the were (B) Examples of persistent crawling (upper panels) and diffusive crawling (lower panels) of *Leptospira* cells obtained by single-cell tracking in 10 s. Colors indicate time courses in the order of red orange yellow green blue, and indigo. (C) Example time courses of leptospires crawling on the kidney cell surfaces; inset, expanded view of cell displacements and schematics of motion patterns. The upper and lower panels show the long-distance migration represented by directive crawling and the limited migration due to frequent reversal, respectively. (D) Examples of MSD vs time plots, evaluating the directivity of individual leptospiral cell movements: Plots with slopes ~ 2 indicate persistently directive crawling (upper), whereas those with ~ 1 indicate a non-directional movement (lower), i.e., motion with high frequency of reversal patterns and short net migration distance (refer to the schematic explanation of motion pattern in insets of **C**). (E) MSD slopes determined for each host-bacterium pair. The boxes show the 25th (the bottom line), 50th (middle), and 75th (top) percentiles, and the vertical bars show the minimum and maximum values. The right panel show the strain-dependence of the medians. The statistical analysis was performed by Mann-Whitney U test (* *P* < 0.05).

## Discussion

Our results suggested the importance of adhesion to and persistent crawling on the host tissue for the pathogenicity of *Leptospira*. The thermodynamic and kinematic parameters are associated in Fig 4A, showing the tendency that pathogenic species are biased to the crawling state and can migrate longer distance on the host tissue surfaces. The crawling motility of *Leptospira* is caused by the attachment of the spirochete cell body to surfaces via adhesive cell surface components(*16, 20*). The successive alternation in the attachment and detachment of adhesins allows for this progressive movement by the spirochetes, however, an excessively strong adhesion can inhibit crawling(*16*). LPS, the molecular basis for the identification of the different *Leptospira* serovars(*2, 12*), is thought to be a crucial adhesin important for this crawling motion(*16*). Thus, it is possible that compatibility of the serological characteristics of leptospires and the surface properties of the host tissue might affect the crawling behavior over the tissue surfaces and the subsequent clinical consequences. The results of our biophysical experiments outline a plausible framework for the adhesion and crawling-dependent pathogenicity of *Leptospira* (Fig. 4B). The biased transition from the adhesion to the crawling state and the long-distance, persistent crawling allows leptospires to explore the host’s cell surfaces, increasing the probability of encountering routes for invasion through their intracellular tight junctions (Fig. 4B, left). In contrast, the swimming or weakly attached leptospires can be swept by external forces, such as intermittent urination, and diffusive crawling by which leptospires cannot be disseminated over host tissues will not be involved in invasion (Fig. 4B, right).

**Fig. 4.**
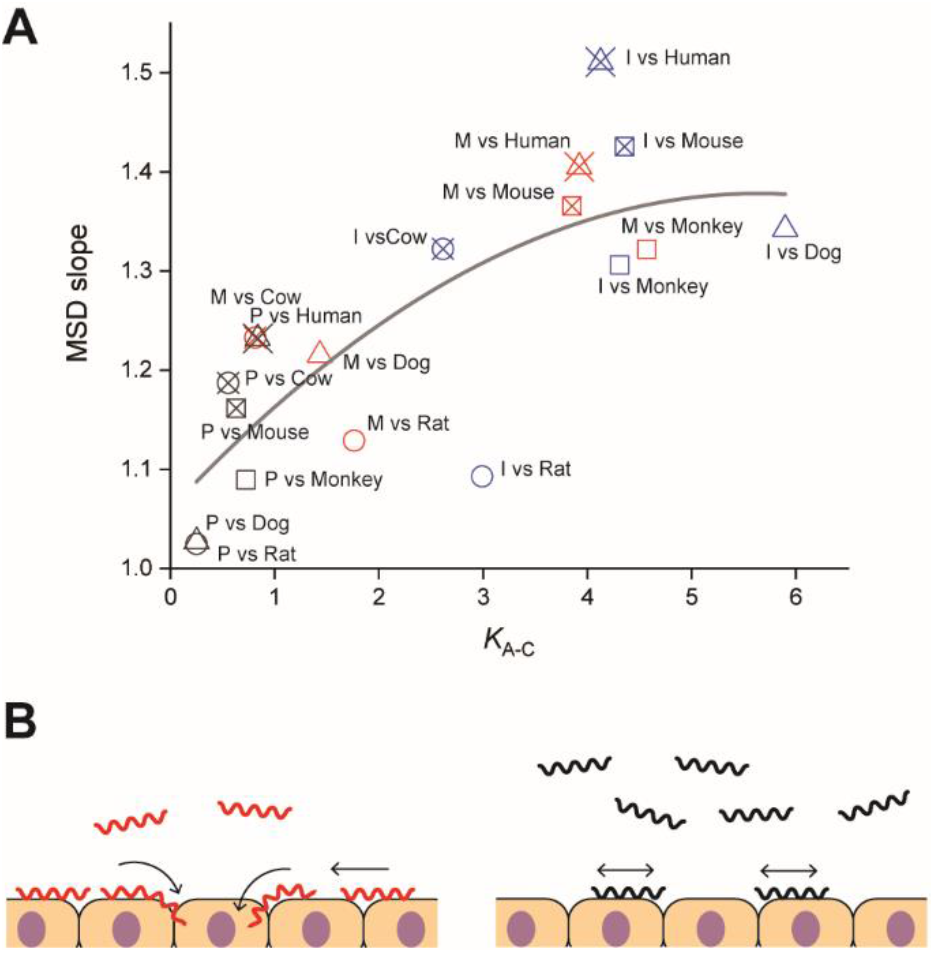
Correlation of adhesion, motility and pathogenicity. (A) Relationship between *K*_A-C_ and MSD slope (i.e., crawling persistency). The gray line is the result of quadratic curve fitting. The correlation coefficient is 0.77. See Fig. 2D for symbols. (B) A plausible model for crawling-dependent pathogenicity of *Leptospira*. In the cases that lead to severe symptoms (left), leptospiral cells were biased to the crawling state, and most of the crawling cells showed directional translation persistently over host tissue surfaces, increasing the invasion probability. For the asymptomatic or non-infectious cases, many leptospires remained in the swimming state and might be removed through body fluids or urination. Some fractions of the adhered leptospires were able to crawl, but their migration distances were limited due to frequent reversal.

Some bacterial pathogens are specialized to invade a very limited array of hosts, whereas others can infect multiple host species. The host range differs for each pathogen and the clinical symptoms depend on each host-pathogen combination. The same applies for leptospirosis, the outcome of *Leptospira* infection depends on the host-serovar association, and some animal species can become an asymptomatic reservoir for particular *Leptospira* serovars. The present experiments also provided data allowing us to discuss the host dependence of the leptospiral dynamics. Among the investigated materials, serovar Manilae vs rats and serovar Icterohaemorrhagiae vs rats are typically asymptomatic pairs, and Fig. 4A shows that the pairs with reservoirs have lower scores in comparison with those causing severe symptoms, such as Manilae vs humans, Icterohaemorrhagiae vs dogs, and others. This implies that the surface dynamics of the spirochete could be related to their host-dependent pathogenicity. Understanding the mechanism of the host preferences by pathogens is important for prevention of the infection spread, but the host-pathogen association in leptospirosis is ambiguous. Also, microarray analyses have revealed that regulation of gene expression in *Leptospira* is affected by its interaction with host cells(*21, 22*). Although host immune responses against *Leptospira* infection remain mostly unknown, pathogenic leptospires interfere with the complement system through the degradation of complement proteins(*23*) and inhibition of coagulation via binding to thrombin(*24*). Thus, abundant factors in both bacteria and hosts should be investigated for a deeper understanding of the host-dependent pathogenicity.

## Materials and Methods

### *Leptospira* strains and growth conditions

Pathogenic serovars of *L. interrogans* serovar Icterohaemorrhagiae (strain WFA135), an isolate from *Rattus norvegicus* in Tokyo, Japan, serovar Manilae (strain UP-MMC-NIID)(*25, 26*) and a saprophytic *L. biflexa* serovar Patoc (strain Patoc I) were used in this study. The serovar of WFA135 was determined by multiple loci variable number of tandem repeats analysis (MLVA) and DNA sequencing of the *lic12008* gene(*26, 27*). Bacteria were cultured in enriched Ellinghausen-McCullough-Johnson-Harris (EMJH) liquid medium (BD Difco, NJ, USA) containing 25 μg/mL spectinomycin at 30 °C for 2 (*L. biflexa*) or 4 (*L. interrogans*) days until the stationary phase. To track *Leptospira* cells when in co-culture with mammalian kidney cells, a green fluorescent protein (GFP) was constitutively expressed in each strain (Fig. 1D; Movie S1).

### Mammalian cells and media

Mammalian kidney epithelial cell lines used included MDCK-NBL2 (dog), NRK-52E (rat), Vero (monkey), MDBK-NBL1 (cow), TCMK-1 (mouse), and HK-2 (human). MDCK, Vero, TCMK, and MDBK cells were maintained in Eagle’s minimum essential media (MEM) (Sigma-Aldrich, Darmstadt, Germany) containing 10% fetal bovine serum or 10% horse serum (Nacalai Tesque, Kyoto, Japan) at 37 °C and 5% CO_2_. NRK cells were maintained in Dulbecco’s modified Eagle’s media (DMEM) (Thermo Fisher Scientific, MA, USA) with 4 mM L-glutamine, adjusted to contain 1.5 g/L sodium bicarbonate and 4.5 g/L glucose, and with 5% bovine calf serum (Nacalai Tesque) at 37 °C and 5% CO_2_. HK-2 cells were maintained in keratinocyte serum-free (KSF) media with 0.05 mg/mL bovine pituitary extract and 5 ng/mL human recombinant epidermal growth factor (Gibco - Thermo Fisher Scientific, MA, USA). All culture conditions contained a 5% antibiotic / antimycotic mixed solution (Nacalai Tesque). Cells were treated with a 0.1% trypsin - EDTA solution (Nacalai Tesque) to dislodge the cells during each passage process.

### GFP expression in *L. interrogans* and *L. biflexa*

For the construction of a replicable plasmid in *L. interrogans*, the corresponding *rep-parB-parA* region of the plasmid pGui1 from the *L. interrogans* serovar Canicola strain Gui44 plasmid pGui1 (*28*) was amplified from a *L. interrogans* serogroup Canicola isolate, and the amplified product was cloned into the PCR-generated pCjSpLe94(*29*) by NEBuilder HiFi DNA Assembly cloning (New England BioLabs), generating pNKLiG1. The *flgC* promoter region and *gfp* were amplified from pCjSpLe94 and pAcGFP1 (Clontech), respectively, and the amplified products were cloned into the *Sal*I-digested pNKLiG1 for *L. interrogans* or the *Sal*I-digested pCjSpLe94 for *L. biflexa*. The plasmids were transformed into strains WFA135, UP-MMC-NIID or Patoc I by conjugation with *E. coli* β2163 harboring the plasmid(*30*). Primer sequences used in this study are listed in Table S1. Expression of GFP did not affect motility in the *Leptospira* serovars (Fig. S3).

### Preparation of kidney cells and *Leptospira* cells in a chamber slide

Kidney cells were harvested with 0.1% trypsin and 0.02% EDTA in a balanced salt solution (Nacalai Tesque) and plated onto a chamber slide (Iwaki, Tokyo, Japan) using their corresponding media without antibiotics. The slides were incubated for 48 h until a monolayer was formed and washed twice with media to remove non-adherent cells. The cells were incubated for a further 2 h at 37 °C and 5% CO2. Approximately 500 ml of stationary phase *Leptospira* cells were harvested by centrifugation at 1,000 *g* for 10 min at room temperature, washed twice in PBS, then resuspended in the corresponding kidney cell culture media without antibiotics at 37 °C to a concentration of 10^7^ cells / ml. These suspensions (1 ml) were then added into the corresponding chamber slides containing the kidney cell layer, and the chamber slides were incubated at 37 °C for 1 h.

### Microscopy observation and adhesion-crawling assay

The movement behaviors, swimming, adhesion and crawling of the *Leptospira* cells on the kidney cells were observed using a dark-field microscope (BX53, Splan 40×, NA 0.75, Olympus, Tokyo, Japan) with an epi-fluorescent system (U-FBNA narrow filter, Olympus) and recorded by a CCD-camera (WAT-910HX, Watec Co., Yamagata, Japan) at 30 frames per second. *Leptospira* cells were tracked using an ImageJ (NIH, MD, USA)-based tracking system and the motion parameters such as motile fraction, velocity and MSD were analyzed using Excel-based VBA (Microsoft, WA, USA). The two-dimensional MSD of individual leptospiral cells during a period Δ*t* was calculated by the following equation: *MSD*(Δ*t*) = 〈(*x*_*i*+Δ*t*_ – *x_i_*)^2^ + (*y*_*i*+Δ*t*_ – *y*)^2^〉, where (*x_i_, y_i_*) is the bacterial position at *I* (see also Supplementary Fig. 1).

### Statistical analysis

All experiments were performed in triplicate. Statistical differences between data were evaluated using a Student’s *t*-test. The data clustering was performed independently for each experiment using the k-means method in OriginPro (OriginLab Corp., MA, USA). The clustering method grouped the data population into a specified number of clusters referring to the Euclidian distance of each data point from the centroid of the cluster, calculated at every clustering process, and reclassifying the data point to the nearest cluster.

## Acknowledgments

We thank Dr. H. Nishimura (Sendai Medical Center) and Dr. C. Toma (University of the Ryukyus) for the generous gift of animal cell lines; and Dr. E. Isogai (Tohoku University) and Dr. H. Yoneyama (Tohoku University) for the experiment reagents and the insightful discussion. This work was supported by the JSPS KAKENHI: 18K07100 for SN and 18J10834 for JX.

## Author contributions

J.X., N.K. and S.N. planned the project; J.X and N.K. carried out the experiments; S.N. set up the optical system and programs for data analysis; J.X. and S.N. analyzed the data; J.X., N.K. and S.N. wrote the paper.

## Competing interests

The authors declare that they have no competing interests.

## Materials & Correspondence

Shuichi Nakamura; Department of Applied Physics, Graduate School of Engineering, Tohoku University, 6-6-05 Aoba, Aoba-ku, Sendai, Miyagi 980-8579, Japan;

## Data availability

The data supporting the findings of this study are available from the corresponding author upon request.

## Supplementary Materials

**Table S1.**
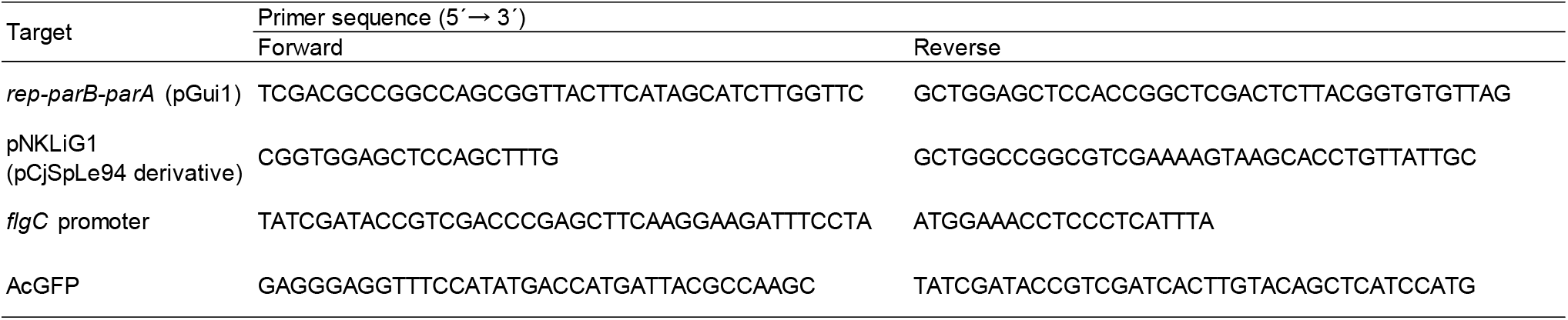
Primer sequences used in this study.

**Fig. S1.**
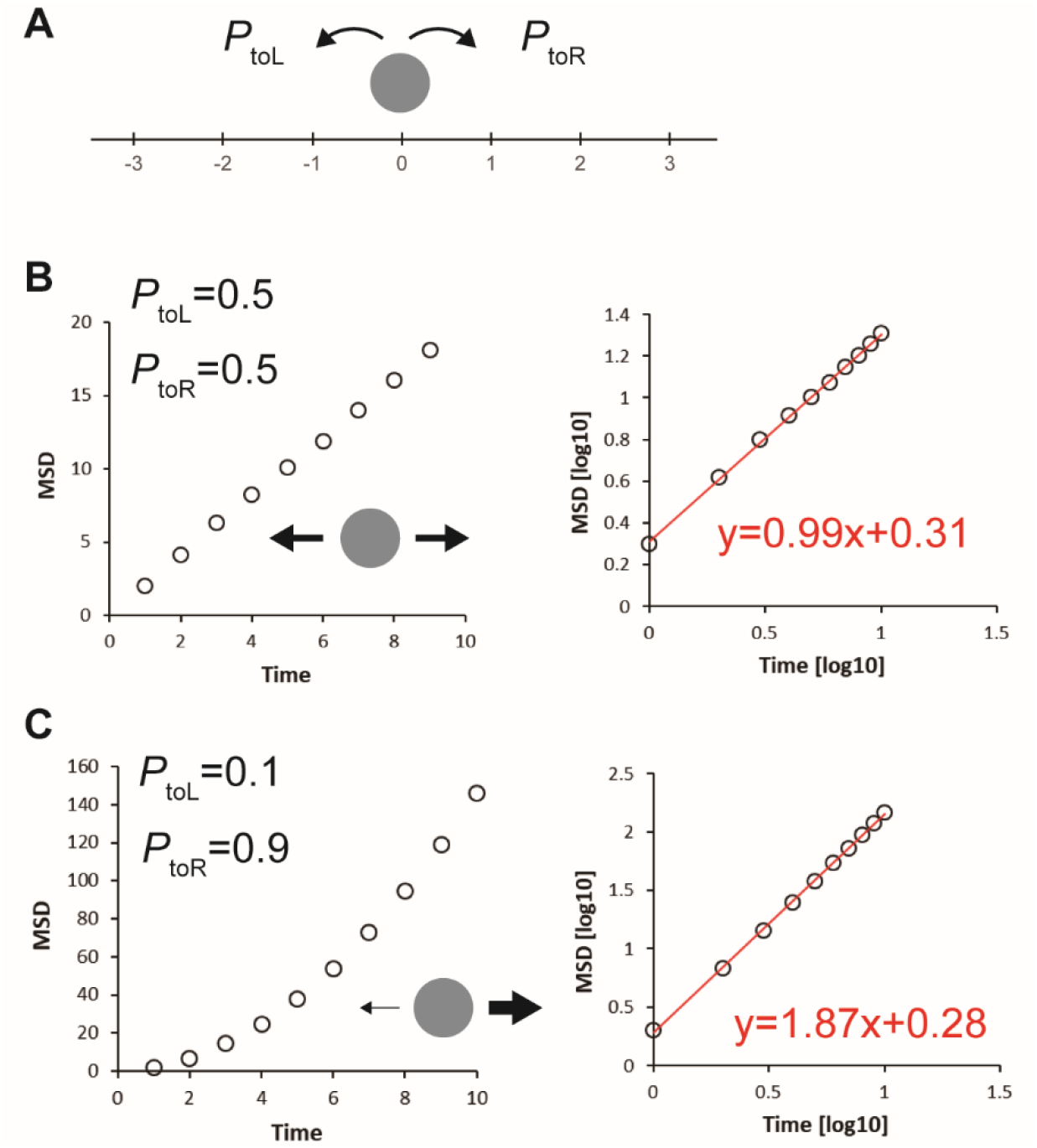
Explanation of MSD plot. (A) To explain the difference in MSD vs time plot of “directed movement” (Figs. 3B-C upper) from that of “diffusive movement” (Figs. 3B-C lower), we performed a computer simulation by considering a particle stepping to the left (−1) or to the right (+1) with the probability of *P*_toL_ and *P*_toR_ (= 1–*P*_toL_), respectively, in a time interval Δ*t*. *P*_toL_ = 0.5 (*P*_toR_ = 0.5) and *P*_toL_ = 0.1 (*P*_toR_ = 0.9) were assumed for simulating simple diffusion and movement biased to the right, respectively, and the step direction was determined by a random number (*rnd*) from 0.0 to 1.0 generated in each event: If *rnd* < *P*_toL_, the particle steps to the left (+1). The time course of the particle position was analyzed as shown in Methods. MSD vs time plots obtained by the simulation show that (B) simple diffusion and (C) directed movement give a linear line and a quadratic curve, respectively (left panels), therefore exhibiting linear lines with slopes of ~ 1 and ~ 2 in double-logarithmic plots (right panels). Red lines are regression lines fitted to data points obtained by simulation.

**Fig. S2.**
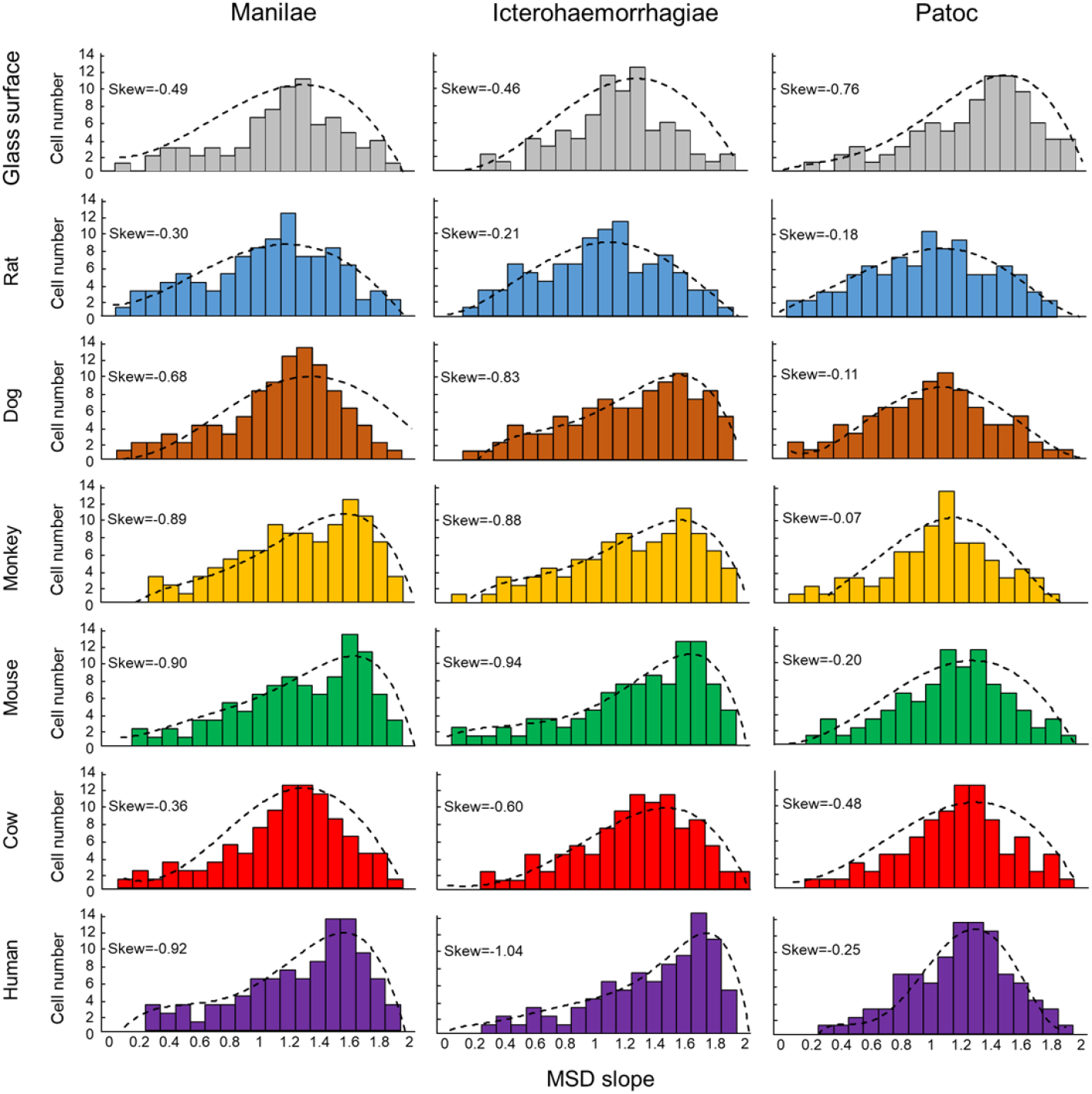
Histograms of the MSD slopes. Dashed lines are the results of curve fitting.

**Fig. S3.**
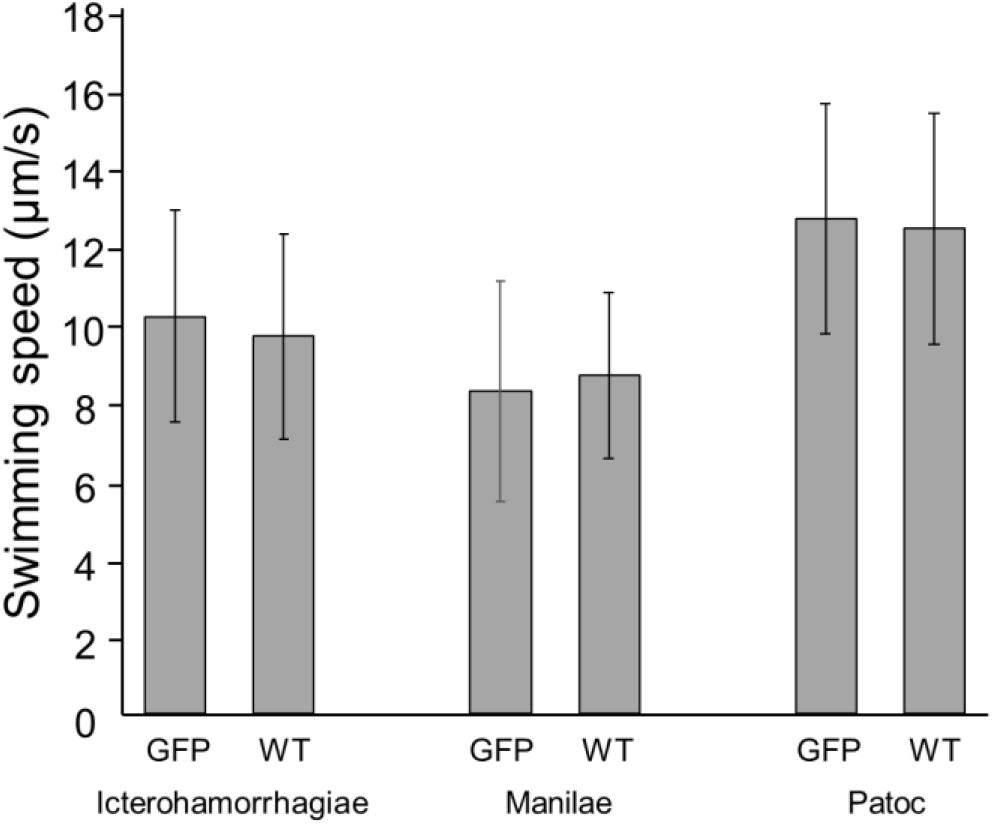
Effect of GFP expression on the *Leptospira* motility. No significant different found between the swimming speeds of GFP expressing strains and wild type strains in each serovar.

### Other Supplementary Materials for this manuscript include the following

Movie S1. Epi-fluorescent images of *L. interrogans* on the rat kidney cell

Movie S2. Progressive, long-distance crawling of *L. interrogans* on the monkey kidney cells

Movie S3. Crawling of *L. interrogans* with highly frequent reversal on the dog kidney cells

## Notes

### Competing Interest Statement

The authors have declared no competing interest.

